# Modeling 3D Mesoscaled Neuronal Complexity through Learning-based Dynamic Morphometric Convolution

**DOI:** 10.1101/2025.08.21.671506

**Authors:** Yik San Cheng, Runkai Zhao, Heng Wang, Hanchuan Peng, Wojciech Chrzanowski, Weidong Cai

**Author notes:** Contributing authors.

## Abstract

Accurate reconstruction of neuronal morphology from three-dimensional (3D) light microscopy is fundamental to neuroscience. Nevertheless, neuronal arbors intrinsically exhibit slender, tortuous geometries with high orientation variability, posing significant challenges for standard 3D convolutions whose static, axis-aligned receptive fields lack adaptability to such complex morphology. To address this, we propose the Dynamic Morph-Aware Convolution (DMAC) framework, which incorporates inherent geometric priors into convolution by jointly adapting both the shape and orientation of the kernel. This enables morphology-aware feature extraction tailored to arborized and variably oriented neuronal trajectories. Specifically, we first apply dynamic tubular convolutions to bridge the structural mismatch between isotropic convolution kernels and the slender morphology of neurons. To sufficiently accommodate the 3D orientation variability of neuronal branches, we further introduce a rotation mechanism that dynamically reorients the tubular kernel via two learnable angles (elevation and azimuth), enabling precise alignment with local neuronal directions. We validate our method through extensive experiments on four mesoscaled neuronal imaging datasets, including two from the BigNeuron project (Drosophila and Mouse) and two additional benchmarks (NeuroFly and CWMBS). Our approach consistently outperforms state-of-the-art methods, achieving average improvements of 5.4% in Entire Structure Average (ESA), 6.9% in Different Structure Average (DSA), and 7.5% in Percentage of Different Structure (PDS). These results demonstrate the effectiveness of our proposed DMAC in capturing complex morphological variations and enhancing structural fidelity across diverse mesoscaled neuronal morphologies.

## 1 Introduction

Accurate reconstruction of neuronal morphology is essential for analyzing and understanding neural circuits and brain network activity. This intricate process traces and digitizes the curved and filamentous neuronal structures from volumetric microscopy images [1–10]. Traditionally, neuron reconstruction has been a labor-intensive and time-consuming process, since it requires cumbersome manual tracing by neuroscientists to annotate intricate and diverse neuronal structures in volumetric images [11]. To advance automated and efficient neuron reconstruction, the DIADEM challenge [12] and BigNeuron challenge [13, 14] were established to benchmark computational methods for large-scale neuronal morphology analysis. More recently, large-scale morphological repositories such as NeuroXiv [15] have enabled open databasing and interactive mining of brain-wide neuronal structures, further facilitating the development and evaluation of automatic reconstruction algorithms. Nevertheless, accurately tracing complex neuronal morphologies remains highly challenging, as the extreme thinness, high curvature, and substantial directional variability of neuronal structures often impede consistent and complete reconstruction.

Traditional automated neuron reconstruction methods predominantly rely on handcrafted algorithms to extract neuronal morphology. Early techniques were broadly categorized into seed-point dependent [16–18] and seed-point independent [19–24] approaches. Seed-point dependent methods initiate tracing from manually or automatically selected points and iteratively extend neuronal branches by following local intensity or connectivity cues. In contrast, seed-point independent algorithms globally search for potential neuronal trajectories. Additionally, feature extraction pipelines based on classical image processing were introduced to enhance weak signals and suppress background noise. For example, Hessian-based filters [25], support vector machine classifiers [26], ray-shooting model [27], and region growing strategies [28] were commonly employed in neuron segmentation and pre-processing. However, many of these tracing algorithms rely on rigid heuristics and hand-tuned parameters. These limit their ability to accurately capture the diverse morphological distribution of neuronal structures, particularly elongated arborization and anisotropically oriented branches.

With the advent of deep learning techniques, data-driven methods have emerged as a powerful alternative to handcrafted approaches, offering improved adaptability to various neuronal morphologies while simultaneously enhancing the quality of raw neuron imaging data. Encoder-decoder based networks, such as 3D U-Net [29] and V-Net [30], are widely used for volumetric neuron segmentation. Various enhancements have been proposed to further improve segmentation performance, including multiscale kernel fusion [31], atrous spatial pyramid pooling [32], graph-based reasoning [33], and knowledge distillation techniques [34, 35]. Moreover, PointNeuron [36] adopts point cloud modeling to capture spatial context and improve geometric representation. Nevertheless, these methods typically lack explicit incorporation of structural priors, limiting their ability to capture complex neuronal structures.

To integrate more structural knowledge, recent works [37–43] have introduced structure-aware modules and geometry-guided learning strategies for neuron segmentation. While effective in improving local structural refinement, they do not explicitly model the intricate morphological characteristics of neurons, such as curvilinear trajectories and directional heterogeneity. Despite these advancements in neuron-specific designs, many segmentation pipelines still adopt popular medical image segmentation frameworks such as nnUNet [44], MedNeXt [45], TransUNet [46], and SegMamba [47]. These models, originally developed for organ or tissue level biomedical segmentation tasks, are mainly optimized for semantic consistency in volumetric data. Nevertheless, these approaches primarily focus on enhancing output-level representations via attention mechanisms, hierarchical feature fusion or auxiliary supervision. They often fall short in adaptively extracting features from structures with high orientation variability and geometric complexity, highlighting the need for more specialized representations tailored to neuronal morphology.

This calls for a feature extraction process that intrinsically encodes morphological priors and dynamically aligns with neuronal geometric variations. As illustrated in Figure 1, neuronal morphology is characterized by extreme thinness, high curvature, and substantial orientation variability within a single 3D volume. Traditional convolutions with fixed, axis-aligned receptive fields inherently lack the flexibility to adapt to such intricate morphological patterns. This often leads to misaligned feature representations, fragmented segmentation, and topological inconsistencies. While recent efforts [48–50] adapt receptive fields for geometric alignment, they lack sufficient directional adaptivity or fail to adjust to local structural variations. To address these limitations, we propose Dynamic Morph-Aware Convolution (DMAC), which equips the model with precise structural adaptation and sufficient 3D directional sensitivity for complex neuronal morphology. DMAC is a 3D convolutional framework that jointly adapts the shape and orientation of kernels to explicitly encode neuronal morphological priors. This enables structure-aware and orientation-sensitive feature extraction, tailored to the filamentous and directionally heterogeneous trajectories of neuronal arbors. Rather than using static, axis-aligned kernels, DMAC reshapes the convolution receptive field into an elongated, deformable tubular form that naturally aligns with the slender geometry of neuronal fibers. To fully capture the anisotropic orientation of neuronal projections, we further introduce a rotation mechanism that dynamically reorients the tubular kernel by two learned orientation angles (elevation and azimuth). The major contributions of this work can be summarized as follows:

- We introduce a novel structure-aware and orientation-adaptive feature extraction paradigm specifically designed for curvilinear neuronal structures. This approach bridges the gap between conventional axis-aligned convolutional kernels and the complex neuronal morphology characterized by extreme slenderness and directional variability.
- We propose the Dynamic Morph-Aware Convolution (DMAC) framework, which enhances morphological learning by reshaping standard 3D convolution kernels into deformable tubular forms and dynamically rotating them via learnable elevation and azimuth angles. This joint structural and directional adaptivity facilitates morphology-aware convolution along filamentous, variably curved neuronal trajectories, thereby improving morphological continuity and topological fidelity in segmentation.
- We conduct extensive experiments on four diverse mesoscale neuron reconstruction datasets, including two from the BigNeuron project [13] (Drosophila and Mouse) and two additional benchmarks (NeuroFly [51] and CWMBS [43]). Our approach consistently demonstrates superior tracing performance over state-of-the-art segmentation methods, yielding an average enhancement of **5.4%** in Entire Structure Average (ESA), **6.9%** in Different Structure Average (DSA), and **7.5%** in Percentage of Different Structure (PDS).

**Fig. 1.**
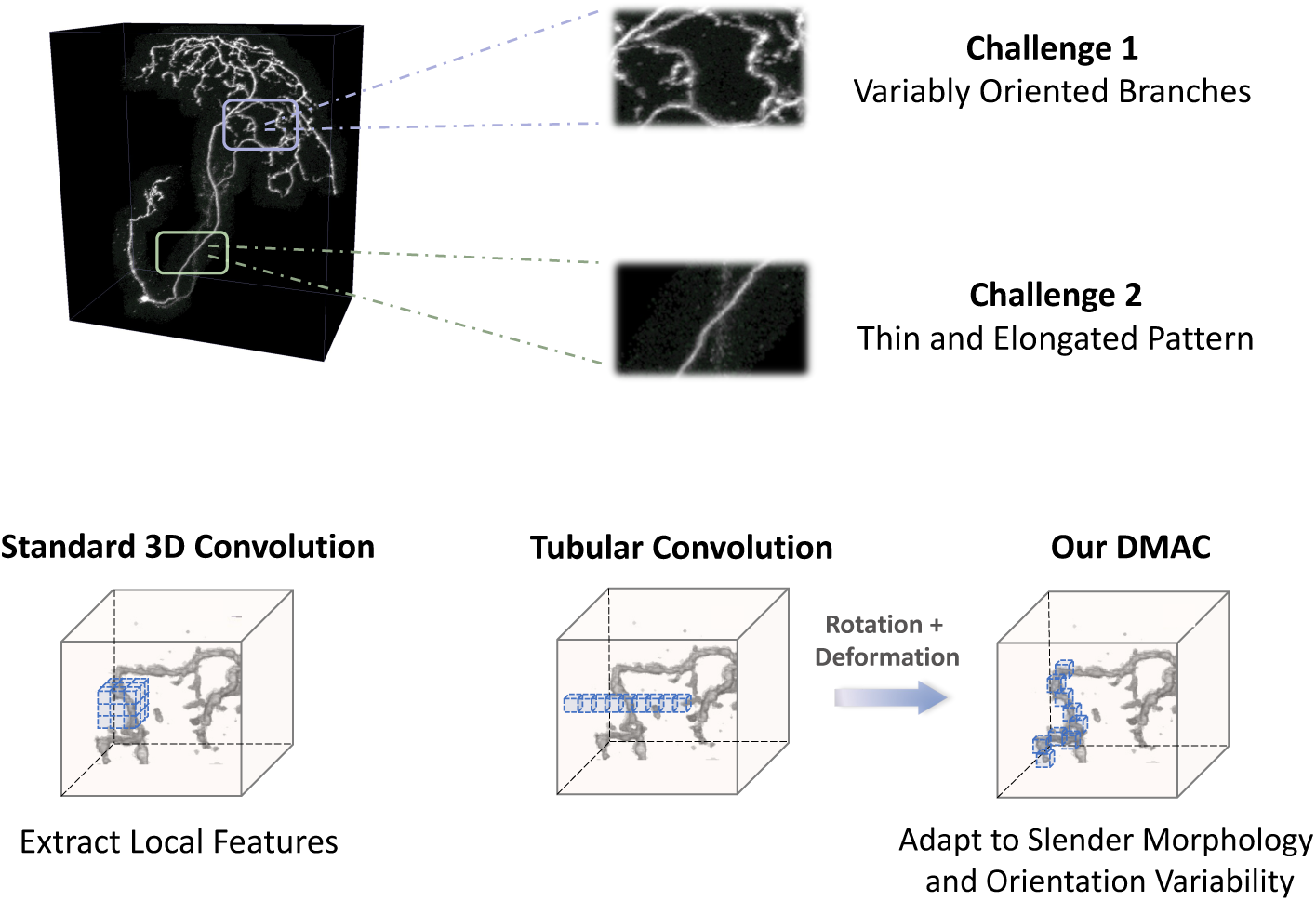
Challenges: Microscopic neuron images often exhibit thin, elongated, and variably oriented structures, as shown in the zoomed-in regions. These complex geometries make it difficult to extract accurate and coherent morphological features. **Motivation:** The standard 3D convolution is designed to extract local features using static, axis-aligned receptive field. Yet, such fixed kernel is insufficient to capture the filamentous and orientation-varying patterns of neurons. This motivates the need for structure-aware and orientation-adaptive convolutional mechanisms.

## 2 Related works

### 2.1 Neuron morphological learning

Traditional automated neuron reconstruction methods primarily rely on handcrafted algorithms to extract neuronal morphology. These include seed-point dependent methods [16–18] that iteratively extend neurites from selected initial points based on local cues, and seed-point independent approaches [19–24] that globally search for neuronal trajectories. Classical pipelines often enhance signals using Hessian filters [25], region growing [28], support vector machine classifiers [26], or ray-shooting model [27]. However, they remain constrained by rigid heuristics and limited adaptivity to morphological variability.

With the rise of deep learning, encoder-decoder architectures such as 3D U-Net [29] and V-Net [30] have demonstrated promise in volumetric neuron segmentation. Subsequent extensions involving multi-scale aggregation [31], graph-based modeling [33], point-cloud representations [36], voxel-wise cross-volume representation learning [52], and unsupervised embedding [53] have further improved performance, while neuron-specific models [37–43] integrate structural priors into network design. Nevertheless, most of these methods focus on output-level refinement and lack a dynamic mechanism to adapt feature extraction process itself to the elongated arborization and anisotropic orientation patterns of neuronal branches.

### 2.2 Dynamic anisotropic convolution designs

To better capture geometric variations, recent efforts [48, 54] have introduced dynamic or deformable convolutions to adapt receptive fields to object geometry. While effective in natural image tasks, these methods remain limited in modeling the elongated and orientation-variant patterns of neuronal branches. To address filamentous geometry more broadly, several works [49, 50] have proposed anisotropic kernel designs which employ elongated convolutions to enhance feature extraction for slender structures. However, these methods lack mechanisms to dynamically adapt to varying directions in neuronal projections. In contrast, adaptive rotated convolution (ARC) [55] dynamically rotates standard convolution kernels to align with object orientation. However, ARC is primarily developed to detect rotated objects in 2D natural images and does not incorporate morphological adaptation, limiting its effectiveness in modeling 3D filamentous neural architectures. Overall, existing dynamic convolutional approaches fall short in jointly adapting both the shape and orientation of receptive fields to align with the directional heterogeneity and filamentous geometry of neuronal trajectories. In contrast, our proposed DMAC framework introduces a unified shape-and-orientation adaptive design, enabling both morphological shaping and directional alignment of kernels. This effectively bridges the gap in modeling slender and orientation-variant neuronal structures.

## 3 Method

We propose the Dynamic Morph-Aware Convolution (DMAC) framework for volumetric neuron segmentation. An overview of the proposed framework is shown in Figure 2. DMAC is designed to explicitly model the filamentous and variably oriented trajectories of neuronal morphology, which pose challenges for standard 3D convolution due to their fixed, axis-aligned receptive fields. To address this, DMAC decomposes the 3D convolution kernel design into three components: (1) Tubular Stretching, which reshapes the receptive field into a tubular form that conforms to elongated neuronal morphology; (2) Dynamic Rotation, which adaptively reorients the tubular kernel to align with local neuronal orientation using two learnable angles; and (3) Cumulative Deformation, which further deforms the kernel to capture fine-grained variations of neuronal structure. Additionally, a multi-view aggregation strategy is used to enrich spatial context by integrating features from multiple anatomical directions. To further improve learning, a deep supervision scheme is utilized to facilitate training and enforce consistent segmentation.

**Fig. 2.**
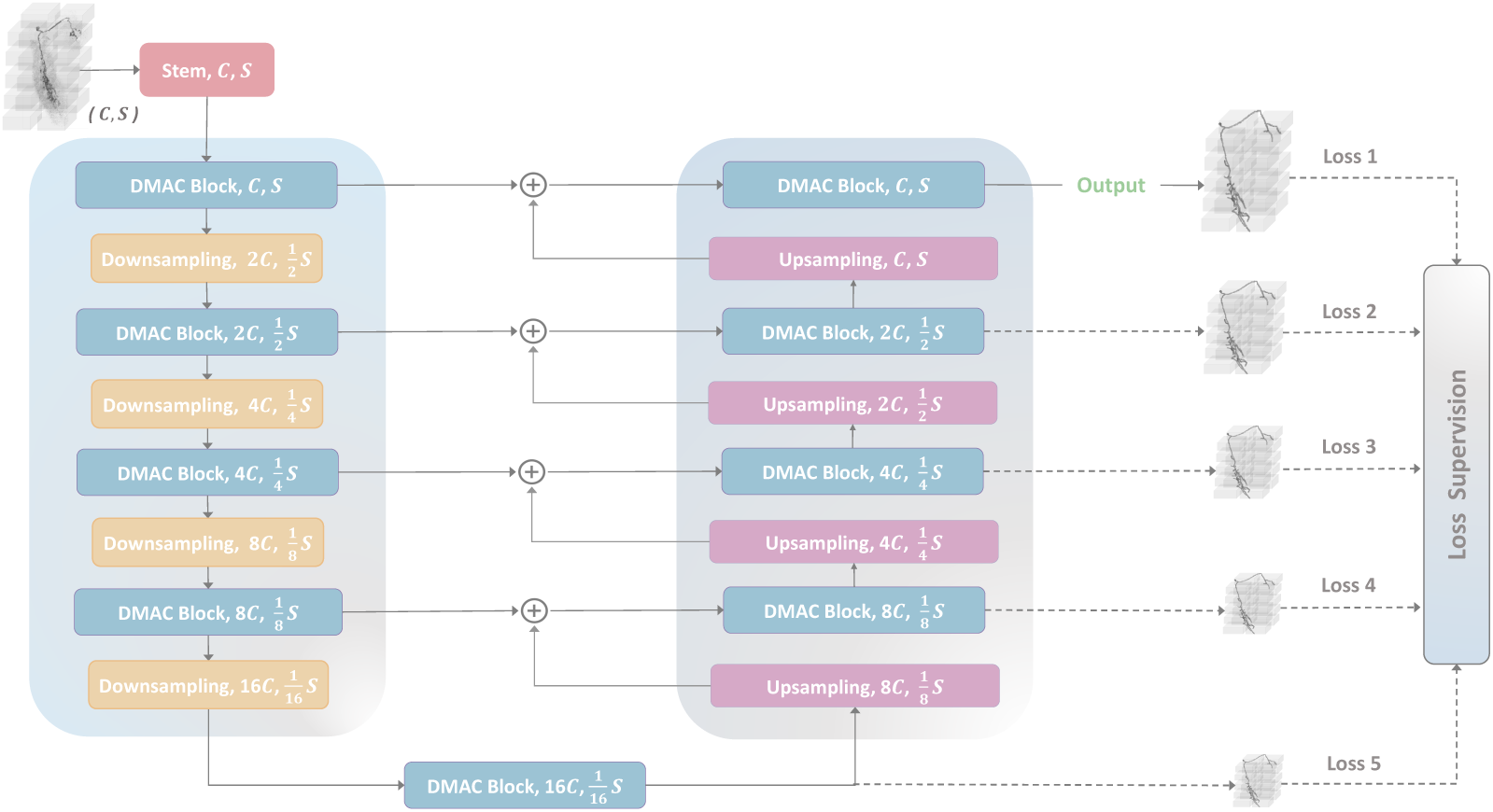
Overview of our framework. Our network adopts a U-shaped encoder-decoder architecture, where Dynamic Morph-Aware Convolution (DMAC) blocks serve as core feature extractors. Downsampling and upsampling modules follow the MedNeXt [45] design to capture multi-scale representations. Skip connections enable cross-scale feature fusion between the encoder and the decoder. Deep supervision is applied at multiple stages (indicated by dashed lines) to facilitate training. *C* denotes the number of feature channels, and *S* denotes the spatial resolution (i.e., input volume shape [*D, H, W*]).

### 3.1 Dynamic morph-aware convolution

Neuronal structures typically exhibit slender, tortuous morphology with varying orientations and sharp directional changes. Traditional 3D convolutions with static, axis-aligned kernels are inherently limited in modeling such intricate geometry. Their fixed receptive fields tend to over-smooth along irrelevant directions while under-capturing critical features aligned with curvilinear neuronal paths, leading to fragmented or inconsistent predictions. To address this, we introduce DMAC that is a novel convolutional design tailored to the unique morphological properties of neurons. Specifically, DMAC stretches the standard 3D convolution into a tubular shape that better conforms to the elongated geometry of neuronal fibers, enabling effective representation along straight or gently curved segments. However, shape adaptation alone lacks directional sensitivity, making it insufficient to capture regions with sharp bends or oblique orientations. We therefore introduce a dynamic rotation mechanism to further reorient the tubular kernel with the dominant local orientation. By jointly modeling the shape and orientation of convolutional receptive field, DMAC achieves structure-aware and orientation-adaptive feature extraction along filamentous and highly curved neuronal trajectories.

#### 3.1.1 Tubular stretching

Traditional 3D convolutions rely on isotropic cubic kernels with receptive fields uniformly distributed across all spatial dimensions. While effective for modeling blob-like structures, such isotropic kernels are inherently suboptimal for extracting the thin, elongated morphology of neuronal fibers. In practice, conventional kernels tend to gather excessive background information while failing to efficiently capture elongated context along neurite trajectories. To better align the kernel with the filamentous neuronal geometry, inspired by [50], we reconstruct the standard 3D convolution into a tubular-shaped kernel that conforms to the morphology of slender neuronal structures. As illustrated in Figure 3, this design stretches the kernel receptive field along a principal axis, enabling more effective context aggregation along filamentous neuronal paths and improving morphology-aware feature extraction. For instance, given a standard 3 × 3 × 3 kernel, we flatten it into a 1 × 1 × 27 tubular kernel with sampling points arranged along, e.g., the *x*-axis. The set of sampling coordinates in a tubular kernel along the *x*-axis can be defined as:

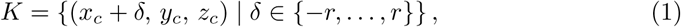

where (*x_c_, y_c_, z_c_*) denotes the coordinate of the center grid, and *δ* represents the distance (in grid units) from the center grid along the *x*-axis. The tubular kernel samples a total of *k* = 2*r* +1 discrete grid positions symmetrically along the *x*-axis with respect to the center grid, forming an elongated receptive field.

**Fig. 3.**
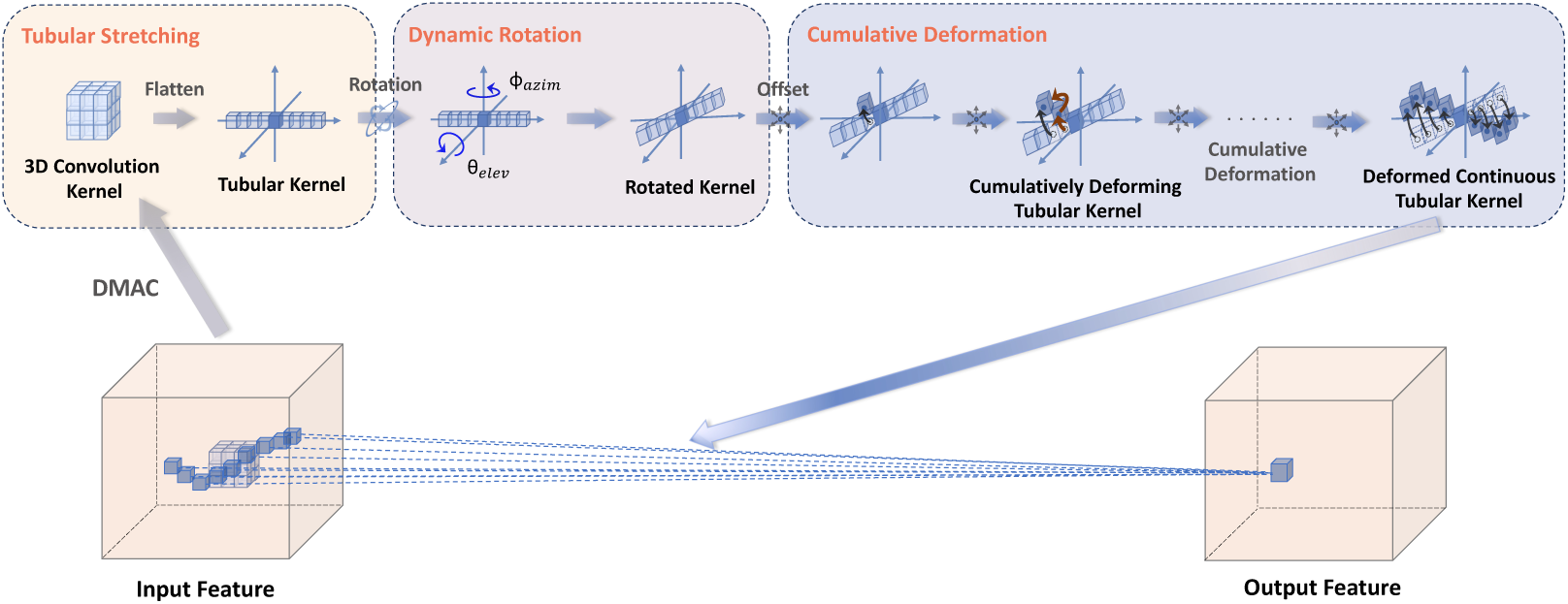
Illustration of the proposed Dynamic Morph-Aware Convolution (DMAC). The input 3D convolutional kernel is first flattened and reshaped into a tubular kernel aligned with a principal axis. This tubular kernel is then dynamically rotated by a pair of predicted angles: elevation angle *θ*_elev_ (tilt in the vertical plane) and azimuth angle *ϕ*_azim_ (rotation in the horizontal plane), aligning the kernel with local neuronal orientations. After rotation, a cumulative offset mechanism progressively deforms the sampling positions along the two non-principal axes, enabling local adaptation to fine-scale morphological variations. The center of the tubular kernel from which deformation is symmetrically accumulated in both directions along the principal axis. The black arrow in the deforming tubular kernel indicates the cumulative offset from the kernel center, while the brown arrows represent the local deformation applied at each sampling step.

Although this morphological adaptation enhances sensitivity to linear neuronal structures, it remains insufficient in regions where neurite trajectories exhibit sharp curvatures or complex directional changes. To address this limitation, we introduce a dynamic rotation mechanism in the following section to enable orientation-aware kernel alignment.

#### 3.1.2 Dynamic rotation

While tubular convolution improves alignment with the elongated shapes of neurons, its fixed sampling axis limits its ability to follow arbitrarily oriented neuron trajectories in 3D space. In practice, neuronal branches often exhibit diverse orientations and oblique curvatures. In regions where neurites twist sharply or extend obliquely, a static tubular kernel fails to adaptively align with the local direction of the neuronal projections, leading to suboptimal feature extraction. To further adapt the tubular kernel to the local structural orientation, we introduce a dynamic rotation mechanism that reorients the tubular kernel along neuronal trajectories and directions via two orientation angles. Specifically, we predict a pair of voxel-wise rotation maps: an elevation map (*θ*) and an azimuth map (*ϕ*). These maps are predicted by:

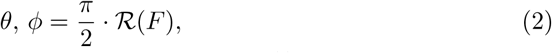

where *F* ∈ R*^C×D×H×W^* is the input feature map, and R(·) denotes a 3D convolutional layer followed by group normalization and softsign activation. The predicted rotation maps *θ, ϕ* ∈ R^1*×*^*^D×H×W^*. At each voxel location, the predicted angles are used to construct a 3D rotation matrix *R* ∈ R^3*×*3^ as the composition of two elementary rotations:

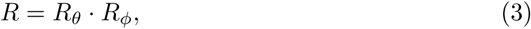

where *R_θ_* and *R_ϕ_* are standard 3D rotation matrices for elevation and azimuthal angles respectively. They are defined as:

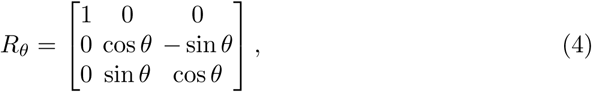

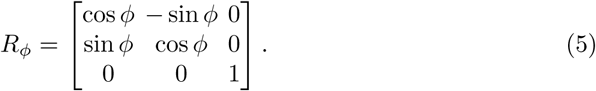

As the tubular kernel slides over the input feature map, the rotation matrix *R* predicted at each voxel location is then applied to rotate all sampling points in the kernel with respect to the kernel center:

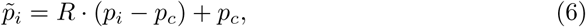

where *p_i_* ∈ R^3^ denotes the original coordinates of the *i*-th sampling point, *p_c_* ∈ R^3^ is the coordinates of kernel center, and *p* ∈*_i_* ∈ R^3^ is the rotated coordinates of the *i*-th sampling point. By rotating the sampling pattern around the kernel center, the tubular kernel effectively adapts to the dominant orientation of local neuronal projection, enabling accurate modeling of curvilinear and variably oriented neuronal structures.

#### 3.1.3 Cumulative deformation

After rotation alignment, we apply a learnable cumulative deformation strategy to capture fine-scale local variations, while preserving structural continuity of the tubular kernel. Unlike conventional deformable convolutions that freely predict independent offsets for each sampling point, leading to spatially disjointed sampling and deviation from slender targets, we adopt a cumulative offset strategy. In this design, the position of each sampling point is recursively determined relative to its predecessor, with all offsets constrained within the range [−1, 1]. This not only enforces continuity but also stabilizes the deformation process, ensuring that the kernel stays aligned with tubular structures. Let the tubular kernel contains *k* sampling points indexed by *i* ∈ {0*, …, k*−1}, with *c* = ⌊*k/*2⌋ denoting the index of the kernel center. The predicted offset field is *O* ∈ R^2^*^k^*, where we predict a 2D offset (Δ*y_i_,* Δ*z_i_*) for each sampling point (assuming the principal axis for the tubular kernel is *x*). Only the coordinates in the two non-principal directions are subject to deformation, while the coordinate along the principal axis (e.g., *x*-axis) remains fixed to preserve tubular consistency and continuity of the kernel. The deformed coordinates of each sampling point are computed by recursively accumulating the incremental offsets outward from the kernel center in both forward (*i > c*) and backward (*i < c*) directions:

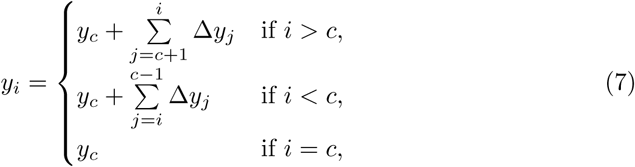

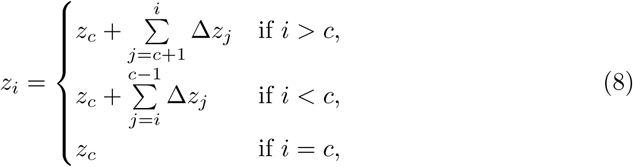

where Δ*y_j_* and Δ*z_j_* are the predicted incremental offsets for the *j*-th sampling point along the y-axis and z-axis, (*y_i_, z_i_*) are the deformed coordinates of the *i*-th sampling point, and (*y_c_, z_c_*) are the coordinates of the kernel center. The coordinate along the principal *x*-axis remains unchanged to maintain the continuity of tubular kernel structure. To prevent abrupt shifts or kernel dispersion, each offset is bounded within the range [−1, 1], ensuring the kernel stays aligned with tubular structures rather than spreading uncontrollably due to large deformations. This process guarantees smooth spatial deformation and simultaneously preserves the tubular shape of the kernel, providing flexibility to adapt to subtle local curvature and structural variations.

#### 3.1.4 Feature interpolation

After cumulative offsets, the final sampling positions typically do not lie on the discrete voxel grids. To obtain meaningful features at these positions, we perform trilinear interpolation over the input feature map, ensuring smooth and differentiable sampling throughout the DMAC module.

### 3.2 Multi-view aggregation strategy

Volumetric neuronal structures are often spatially complex. Using a single-axis tubular kernel is thus insufficient to fully capture their geometric features from multiple anatomical views. To address this, we employ a multi-view learning strategy as shown in Figure 4, in which three tubular kernels, each aligned along one of the principal axes, are applied in parallel:

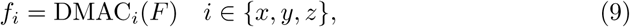

where *F* denotes the input feature map, and DMAC*_i_* applies the tubular kernel along the *i*-axis. The resulting features are then aggregated to form a more comprehensive representation:

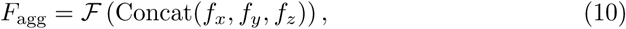

where F(·) denotes a fusion operator, implemented as a 1 × 1 × 1 convolution operation followed by normalization and activation. This multi-view aggregation enables the network to integrate directional features from multiple anatomical perspectives, improving its robustness to direction variance in neuronal morphology.

**Fig. 4.**
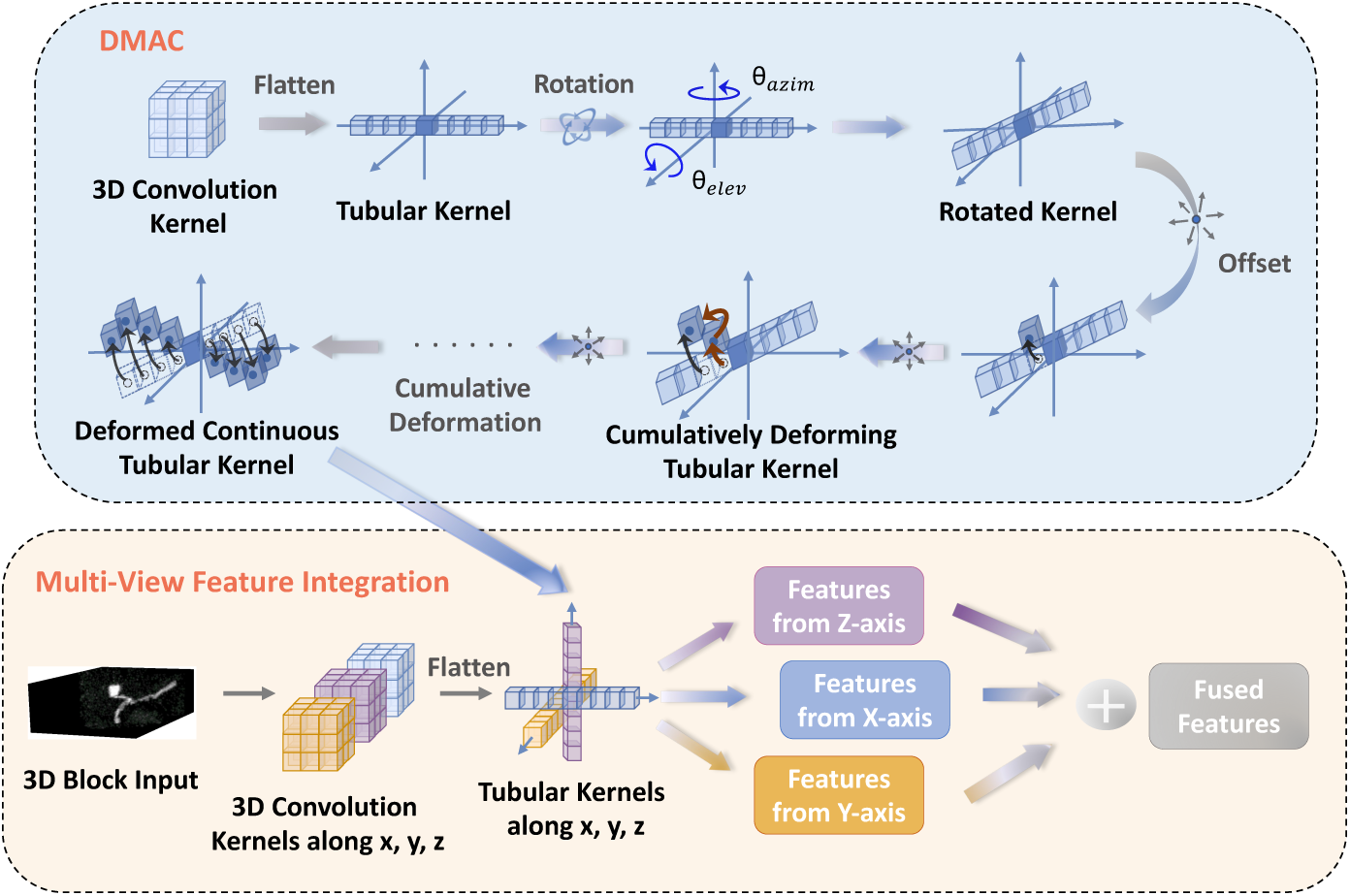
Illustration of multi-view feature integration. Given a 3D input block, three standard 3D convolutions are extended into tubular kernels along the orthogonal axes (*x*, *y*, and *z*). Each tubular kernel extracts axis-specific features, which are further aligned via dynamic rotation and deformation before being fused into a unified representation.

### 3.3 Deep supervision and loss functions

Following common practice in medical segmentation [44], we apply deep supervision to enhance gradient propagation and enforce multi-scale feature refinement. This technique ensures segmentation consistency across multi-resolution levels, resulting in a more robust and coherent neuronal segmentation. The final training objective is computed as a weighted sum of losses from multiple scales, where each output branch is supervised using a combination of Dice loss and Cross-Entropy (CE) loss.

#### Dice loss

The Dice is a loss function that measures the overlap between prediction and ground truth. It is defined as:

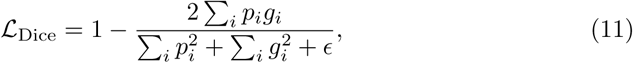

where *p_i_*and *g_i_* represent the predicted probability and ground-truth label at voxel *i*, and *ɛ* is a small constant used to prevent division by zero.

#### Cross-entropy loss

The Cross-Entropy loss is employed to ensure voxel-wise classification accuracy:

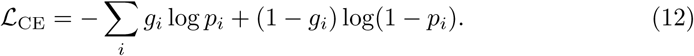

Total loss function is formulated as:

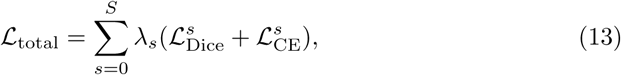

where *S* represents the number of deep supervision outputs, *λ_s_* is a weighting factor for each scale, and 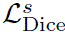 and 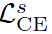 denote the Dice loss and Cross-Entropy loss computed at scale *s* respectively.

## 4 Experiments and results

We evaluate the proposed DMAC framework on four public benchmarks, including two datasets from the BigNeuron project [13] (Drosophila and Mouse) as well as the NeuroFly [51] and CWMBS [43] datasets. These benchmarks represent mesoscale neuronal imaging data, capturing large 3D volumes with intricate arborized structures and significant morphological diversity. Together, these benchmarks cover a wide spectrum of neuronal morphologies, imaging modalities, and signal qualities, providing a comprehensive testbed for 3D neuron segmentation and reconstruction. Comparisons with state-of-the-art methods demonstrate the superiority of our approach in capturing complex neuronal morphology. In addition, we perform an ablation study to analyze the contribution of the key components within the DMAC framework.

### 4.1 Datasets

We conduct experiments on four publicly available volumetric neuron datasets, including two standard benchmarks from the BigNeuron project [13], and two datasets from NeuroFly [51] and CWMBS [43]. All datasets consist of grayscale 3D image stacks acquired by high-resolution light microscopy and paired with expert annotations. These datasets represent mesoscale brain imaging volumes with complex, arborized neuronal structures.

- **BigNeuron** [13]: This benchmark comprises diverse neuron image stacks acquired from various species and imaging settings. In our study, we utilize two representative subsets: the *Drosophila*, which contains 42 volumetric microscopy images of drosophila neurons with varied anatomical features (4 volumes for testing); and the *Mouse*, consisting of 22 high-resolution volumetric images of mouse neurons (3 volumes for testing), generating morphological maps of individual neurons across the entire mouse brain.
- **NeuroFly** [51]: This dataset is constructed from a collection of high-resolution neuron images acquired using VISoR [56] and fMOST [57] imaging techniques, covering both macaque and mouse brain samples. We use 91 volumes for training and 62 volumes for testing. The dataset features complex arborization, varying signal quality, and high morphological diversity.
- **CWMBS (Complex Whole Mouse Brain Sub-Image)** [43]: This dataset consists of 245 volumetric blocks of mouse brain. It contains two categories of image quality: 83 images with strong background noise (49 volumes for training, 34 volumes for testing) and 162 images with weak fluorescence signals (97 volumes for training, 65 volumes for testing).

### 4.2 Performance measures

To comprehensively assess the performance of our proposed method against other baselines, we quantitatively evaluate their segmentation results and their corresponding reconstruction performance. Specifically, we use four metrics for segmentation and three for tracing, all of which are widely adopted in 3D neuron analysis.

#### 4.2.1 Segmentation metrics

We evaluate segmentation accuracy using Precision, Recall, F1 Score, and the 95th percentile Hausdorff Distance (HD95). The first three metrics are defined as follows:

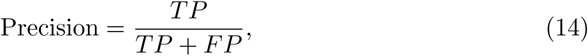

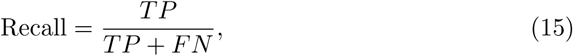

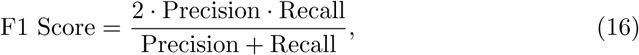

where *TP* (True Positive) is the number of voxels correctly predicted as foreground, *FP* (False Positive) is the number of background voxels incorrectly predicted as foreground, and *FN* (False Negative) is the number of foreground voxels that are incorrectly predicted as background. Precision measures the proportion of correctly predicted foreground voxels among all voxels predicted as foreground. Recall quantifies the proportion of actual foreground voxels that are correctly predicted. For all of these metrics, higher values indicate better segmentation performance. HD95 computes the 95th percentile of the Hausdorff Distance between prediction and ground truth, reflecting surface deviation while discounting outliers. A lower HD95 indicates better geometric alignment between predicted and reference shapes.

#### 4.2.2 Reconstruction metrics

To assess reconstruction quality, we adopt three widely used structure-level metrics:

- Entire Structure Average (ESA): the average minimum distance from every node in the predicted reconstruction to its closest node in the ground truth.
- Different Structure Average (DSA): the average distance of non-overlapping regions between prediction and ground truth.
- Percentage of Different Structure (PDS): the percentage of mismatched branches relative to total length.

For all these tracing metrics, lower values indicate improved topological alignment and reconstruction fidelity.

### 4.3 Data preprocessing and augmentation

To ensure robust training and improve generalization, we apply a series of preprocessing and augmentation steps to all volumetric data:

- Resampling: The raw images exhibit anisotropic voxel spacing. We resample them to 1.0 mm isotropic spacing.
- Intensity Normalization: To reduce intensity variations across images, we normalize voxel intensities to zero mean and unit variance.
- Patch-Based Cropping: Due to GPU memory constraints, we adopt a patch-based training strategy by randomly extracting 3D patches of size 32 × 32 × 32.
- Data Augmentation: To mitigate overfitting caused by limited training data, we apply extensive augmentations including random rotation, flipping, scaling, cropping, elastic deformation, gamma correction, brightness/contrast jittering, and Gaussian noise injection.

### 4.4 Hardware and implementation details

All experiments are conducted on two NVIDIA RTX 4060 GPUs (16 GB VRAM for each). Our networks are trained for 110 epochs using the AdamW optimizer. The learning rate is initialized to 0.001. We employ a combination of Dice loss and Cross-Entropy loss as the loss function. In addition, we use automatic mixed precision (AMP) to accelerate training while reducing memory usage. During inference, a sliding-window approach is adopted to predict testing images, where the window size equals the patch size used in training.

### 4.5 Results

#### 4.5.1 Evaluation on subsets from the BigNeuron project

We evaluate our method on two subsets from the BigNeuron project: Drosophila and Mouse. Their segmentation results are shown in Figure 5 and Figure 7. Our method achieves the highest F1 score on both datasets (49.04% for Drosophila and 53.82% for Mouse) and the lowest HD95 (3.02 and 7.90, respectively). Compared with the strongest-performing baseline, our approach improves F1 by 1.47 percentage points on Drosophila and 1.77 percentage points on Mouse, while reducing HD95 by 0.11 and 1.48, respectively. These improvements demonstrate the robustness of the proposed morphology-aware convolutional design across diverse neuronal morphologies.

**Fig. 5.**
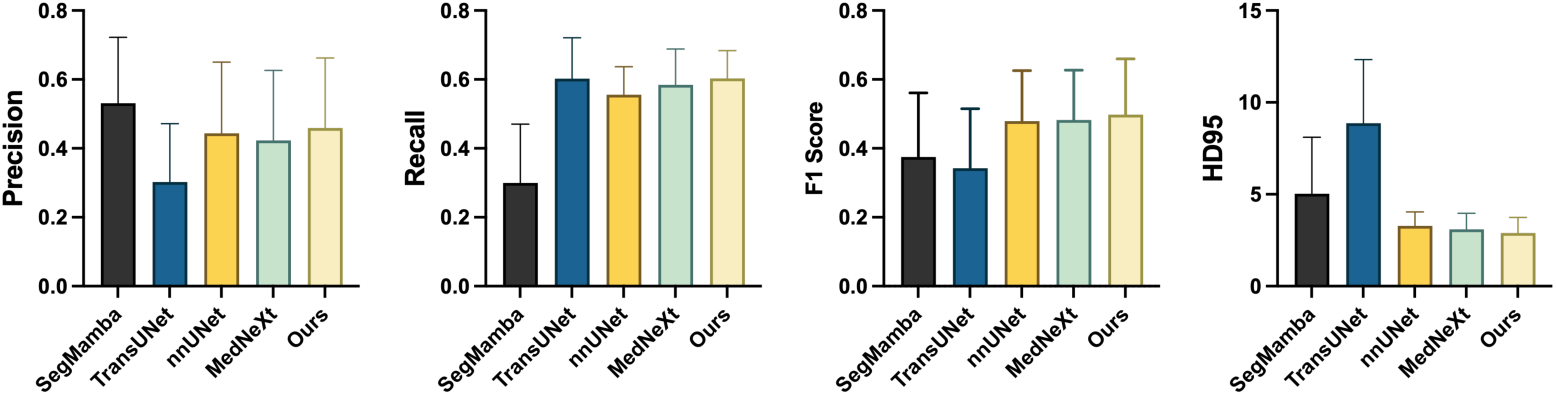
Quantitative segmentation performance on the Drosophila dataset. Bar plots show the mean and standard deviation for Precision, Recall, F1 score, and 95th percentile Hausdorff distance (HD95) across test volumes, comparing SegMamba, TransUNet, nnUNet, MedNeXt, and our proposed method. Higher values of Precision, Recall, and F1 score, and lower values of HD95 indicate better performance.

**Fig. 6.**
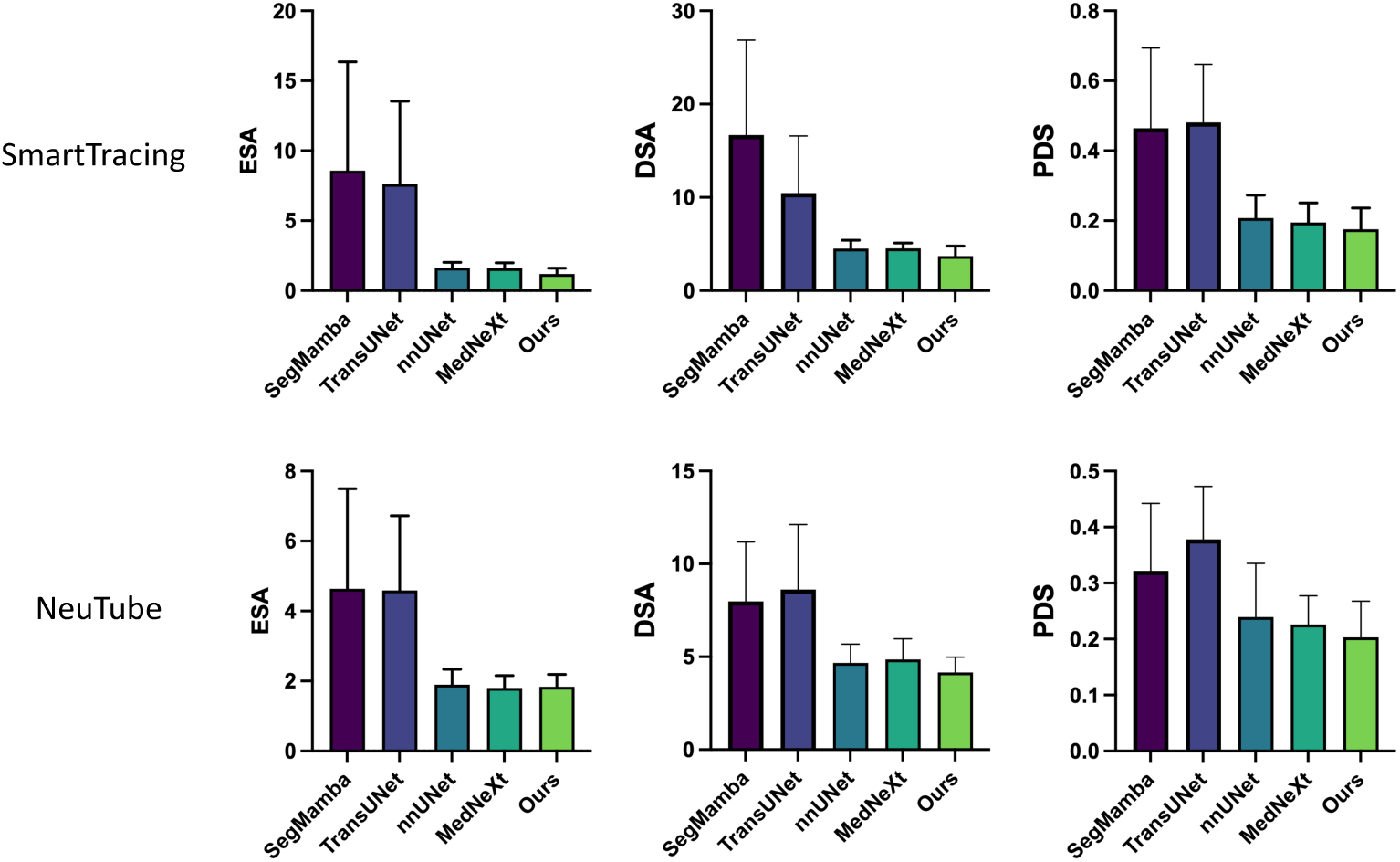
Quantitative neuron reconstruction performance on the Drosophila dataset using two tracing methods: SmartTracing (top row) and NeuTube (bottom row). Bar plots present the mean and standard deviation of ESA, DSA, and PDS across test volumes, comparing SegMamba, TransUNet, nnUNet, MedNeXt, and our proposed method. Lower values indicate better performance.

**Fig. 7.**
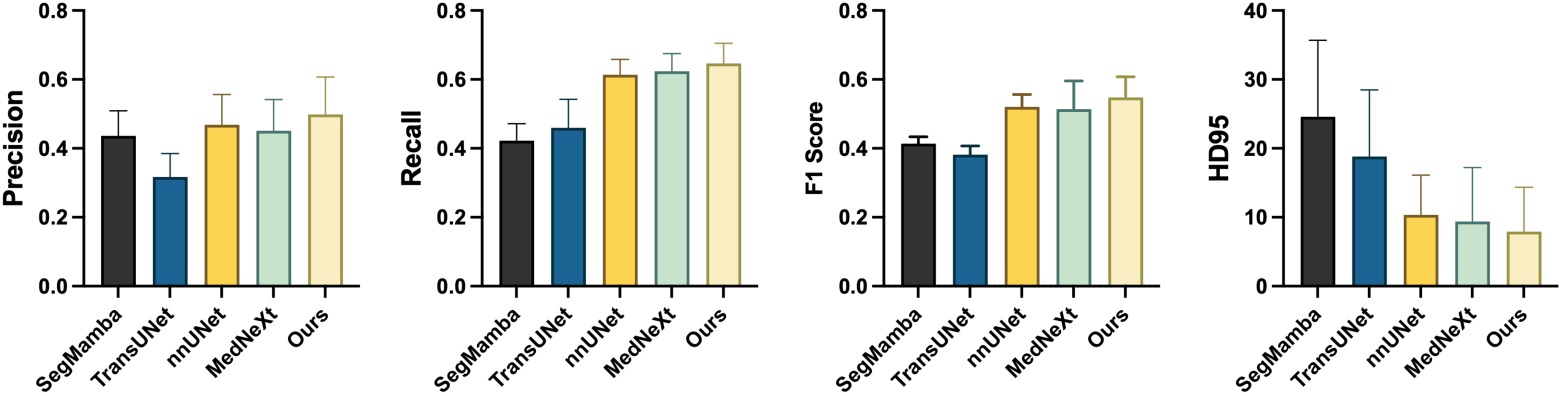
Quantitative segmentation performance on the Mouse dataset. Bar plots show the mean and standard deviation for Precision, Recall, F1 score, and 95th percentile Hausdorff distance (HD95) across test volumes, comparing SegMamba, TransUNet, nnUNet, MedNeXt, and our proposed method. Higher values of Precision, Recall, and F1 score, and lower values of HD95 indicate better performance.

By comparing local details of the segmented outcomes across different baselines, Figure 17 and Figure 18 further corroborate these quantitative improvements.

The zoomed-in views of blue-boxed regions highlight areas where baseline methods exhibit fragmented, discontinuous, or topologically inconsistent predictions. In contrast, our method reduces segmentation discontinuities and enhances topological coherence, resulting in improved overall morphology preservation and fewer fragmented structures.

Tracing results for both datasets, obtained using SmartTracing and NeuTube, are presented in Figure 6 and Figure 8. Our method achieves consistent gains across ESA, DSA, and PDS metrics on both datasets, indicating superior completeness and consistency of the reconstructed neuronal morphology. Qualitative comparisons (Figure 13 and Figure 14) further reveal that our reconstructions are more continuous and morphologically faithful, with fewer disconnections and better alignment to ground truth.

**Fig. 8.**
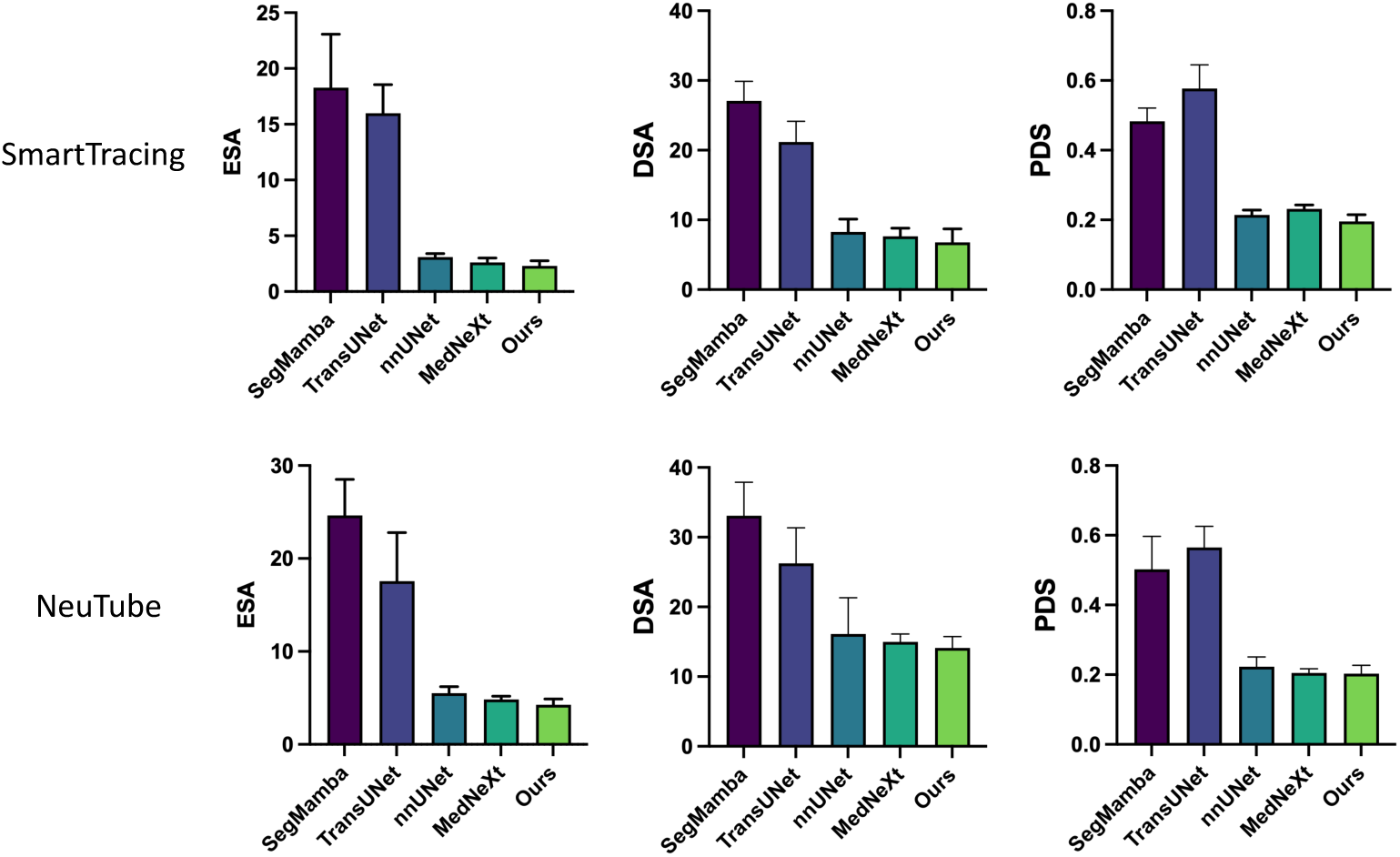
Quantitative neuron reconstruction performance on the Mouse dataset using two tracing methods: SmartTracing (top row) and NeuTube (bottom row). Bar plots present the mean and standard deviation of ESA, DSA, and PDS across test volumes, comparing SegMamba, TransUNet, nnUNet, MedNeXt, and our proposed method. Lower values indicate better performance.

#### 4.5.2 Evaluation on the NeuroFly dataset

Segmentation results on the NeuroFly dataset are presented in Figure 9. Our proposed DMAC-based framework achieves the highest F1 score (64.95%) and the lowest HD95 (16.74), outperforming the second-best method by 1.25 percentage points in F1 and reducing HD95 by 1.46. These results indicate that our morphology-aware design effectively enhances segmentation accuracy and robustness, even in anatomically challenging regions.

**Fig. 9.**
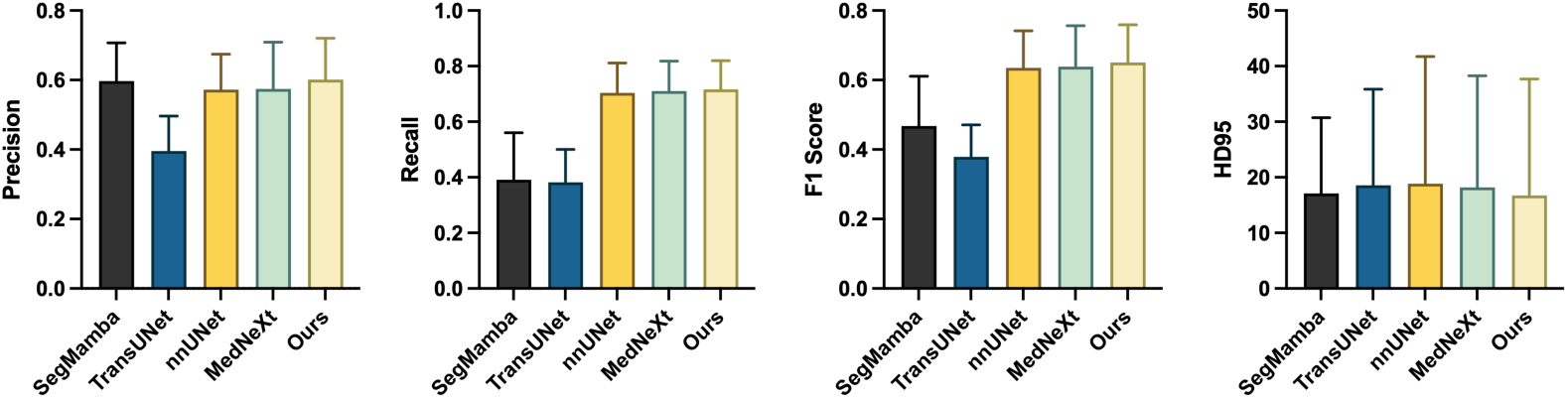
Quantitative segmentation performance on the NeuroFly dataset. Bar plots show the mean and standard deviation for Precision, Recall, F1 score, and 95th percentile Hausdorff distance (HD95) across test volumes, comparing SegMamba, TransUNet, nnUNet, MedNeXt, and our proposed method. Higher values of Precision, Recall, and F1 score, and lower values of HD95 indicate better performance.

The reconstruction performance, evaluated with SmartTracing and NeuTube, is shown in Figure 10. Our method consistently achieves superior results across all tracing metrics, including ESA, DSA, and PDS, demonstrating improved structural continuity and alignment to the ground truth. Qualitative comparisons in Figure 15 further reveal that our reconstructions exhibit fewer fragmented branches and better morphological preservation compared to competing methods.

**Fig. 10.**
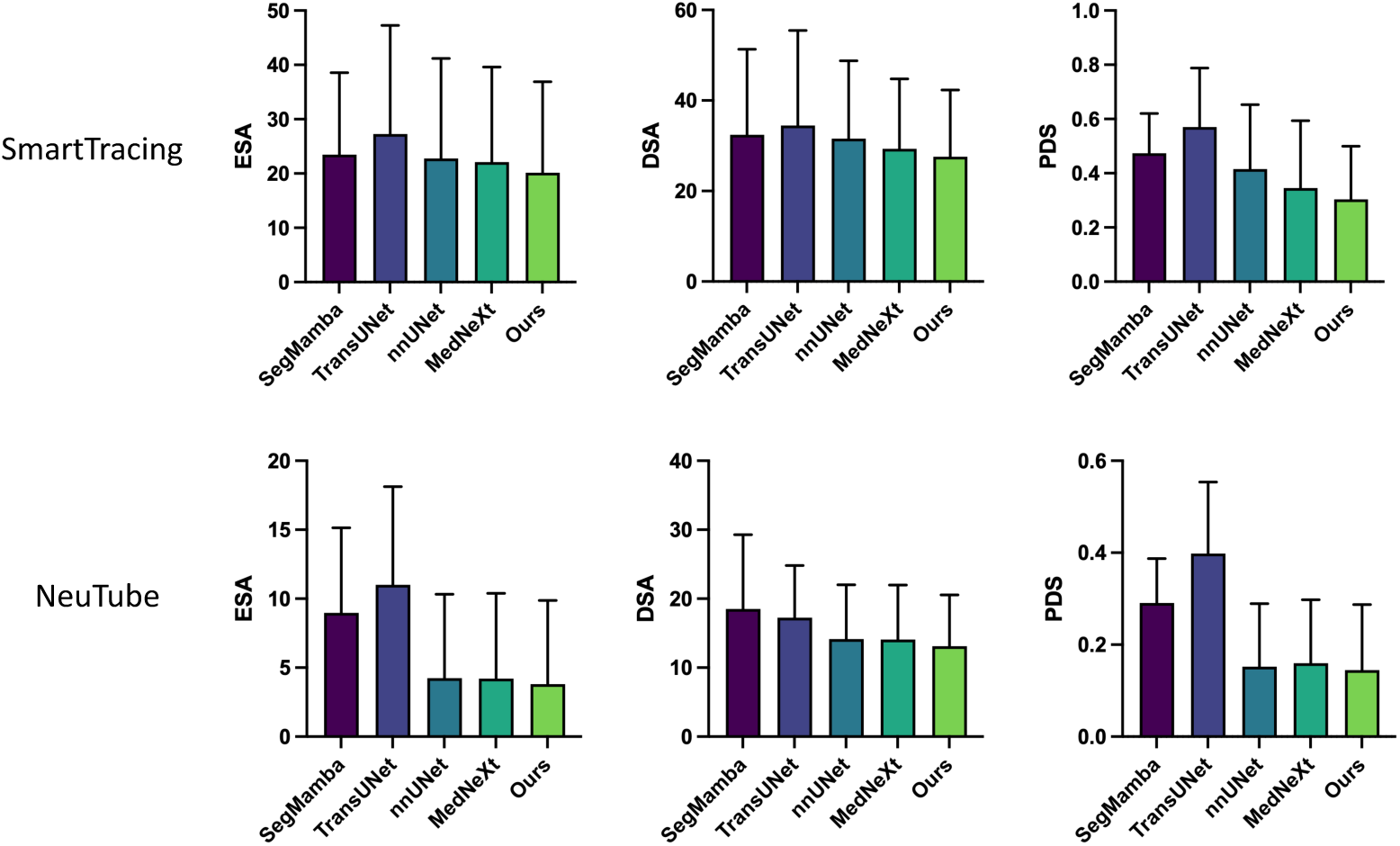
Quantitative neuron reconstruction performance on the NeuroFly dataset using two tracing methods: SmartTracing (top row) and NeuTube (bottom row). Bar plots present the mean and standard deviation of ESA, DSA, and PDS across test volumes, comparing SegMamba, TransUNet, nnUNet, MedNeXt, and our proposed method. Lower values indicate better performance.

#### 4.5.3 Evaluation on the CWMBS dataset

The segmentation performance on the CWMBS dataset is shown in Figure 11. Our method obtains the highest F1 score (37.29%) and the lowest HD95 (14.52) among all compared baselines, surpassing the second-best method by 0.79 percentage points in F1 and reducing HD95 by 1.82. While the numerical gain in F1 is modest, it is accompanied by a clear reduction in HD95, suggesting improved accuracy in delineating fine neuronal structures.

**Fig. 11.**
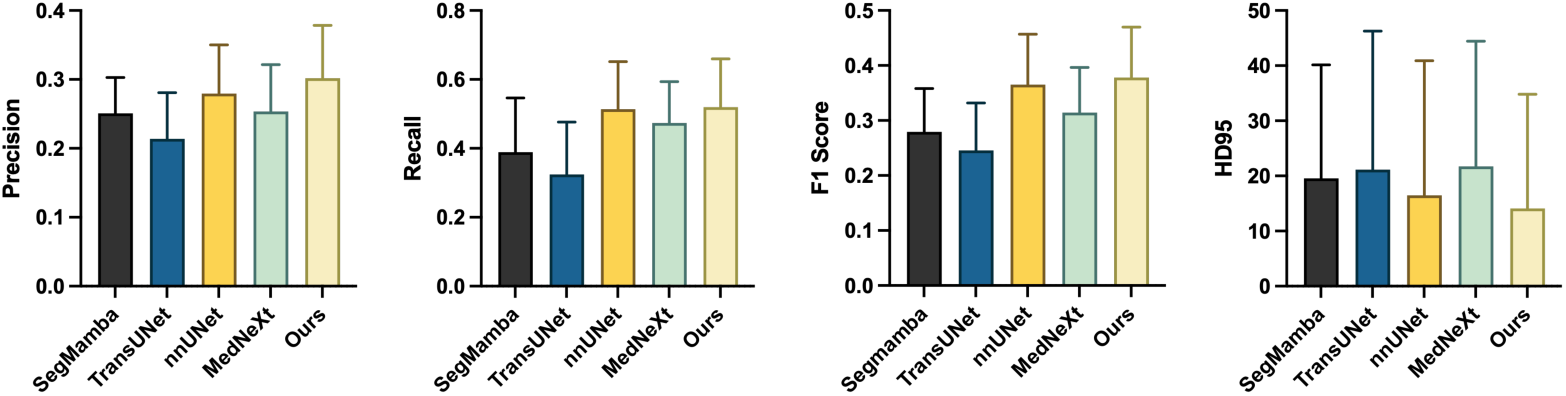
Quantitative segmentation performance on the CWMBS dataset. Bar plots show the mean and standard deviation for Precision, Recall, F1 score, and 95th percentile Hausdorff distance (HD95) across test volumes, comparing SegMamba, TransUNet, nnUNet, MedNeXt, and our proposed method. Higher values of Precision, Recall, and F1 score, and lower values of HD95 indicate better performance.

Tracing results in Figure 12 further confirm this advantage. Under both SmartTracing and NeuTube pipelines, our method achieves the best ESA, DSA, and PDS values, indicating more complete and morphologically faithful neuron reconstructions. Qualitative comparisons are shown in Figure 16, where two representative 3D volumes from different imaging conditions are visualized. Compared with baseline methods, our reconstructions show greater continuity and structural integrity, with fewer disconnections and better alignment to neuronal morphology. These visual results further support the robustness of our method in handling different image conditions and maintaining topological consistency across varying signal qualities.

**Fig. 12.**
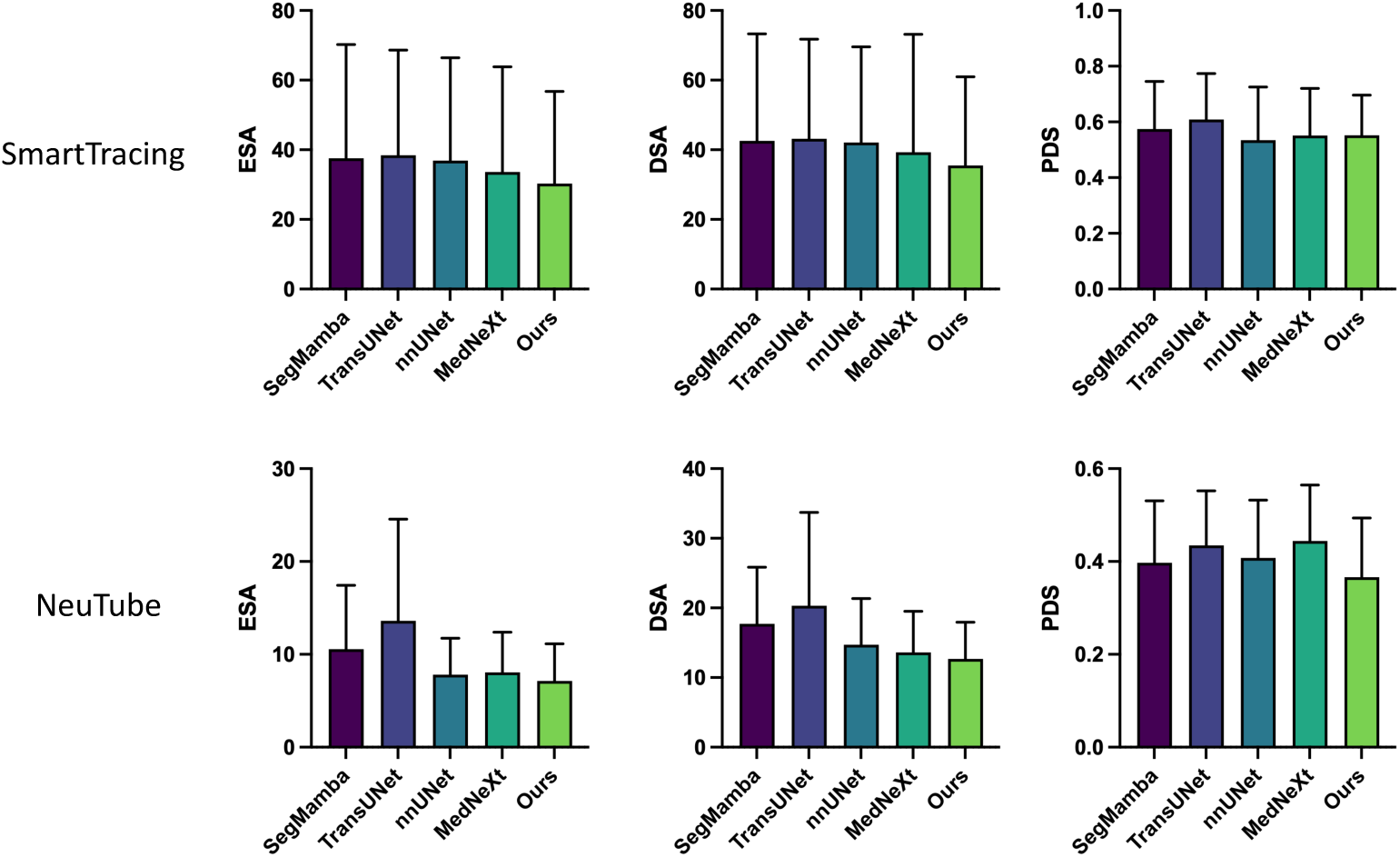
Quantitative neuron reconstruction performance on the CWMBS dataset using two tracing methods: SmartTracing (top row) and NeuTube (bottom row). Bar plots present the mean and standard deviation of ESA, DSA, and PDS across test volumes, comparing SegMamba, TransUNet, nnUNet, MedNeXt, and our proposed method. Lower values indicate better performance.

**Fig. 13.**
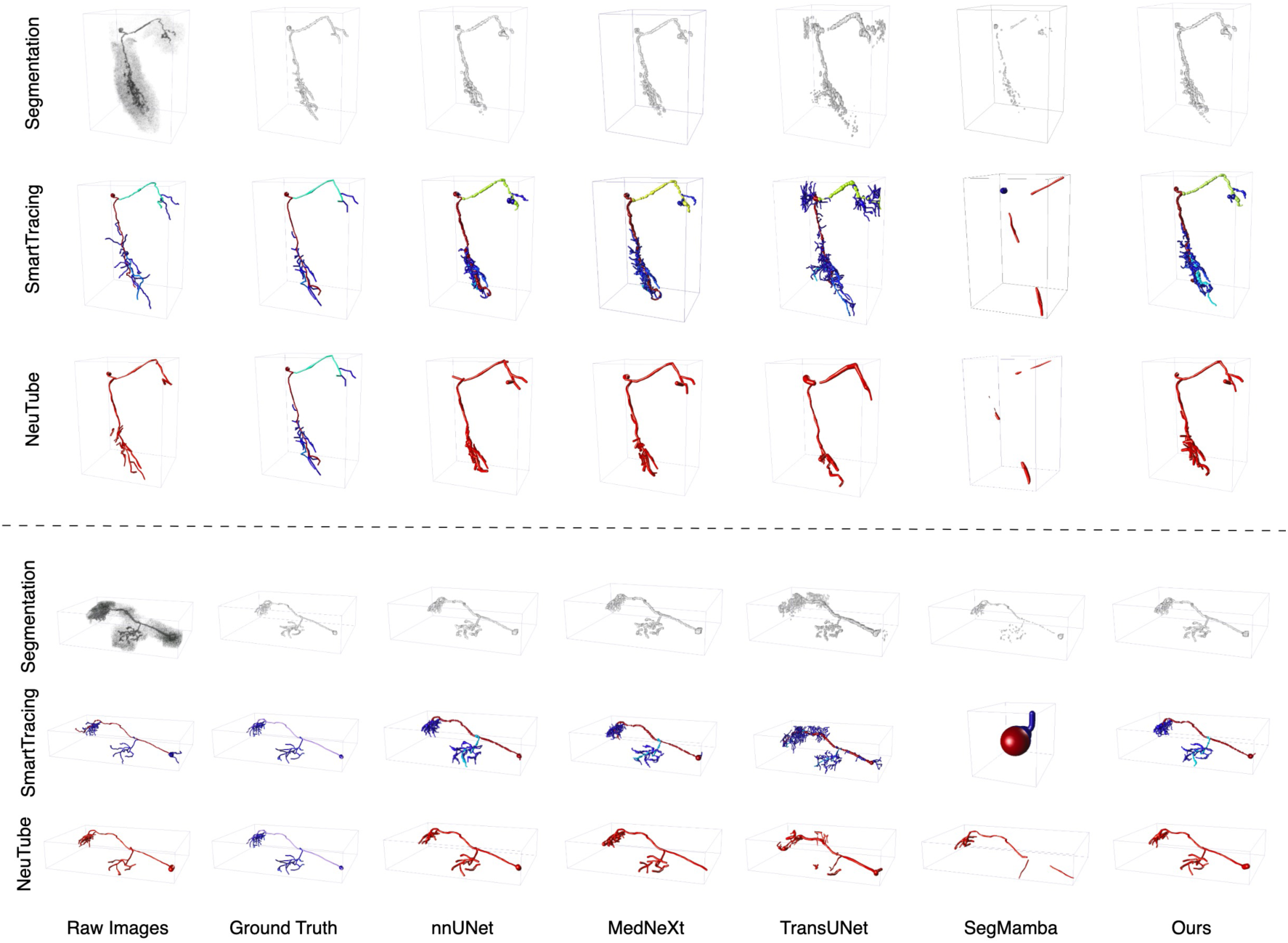
Visualization comparison of the neuron segmentation and tracing performance across different models on the Drosophila dataset. The figure shows two representative cases. For each case, the top row presents the raw image and segmentation outputs from different models, followed by two rows showing the corresponding neuron tracing results generated by SmartTracing and NeuTube. Zoom-in views highlighting fine-grained segmentation details are shown in Figure 17.

**Fig. 14.**
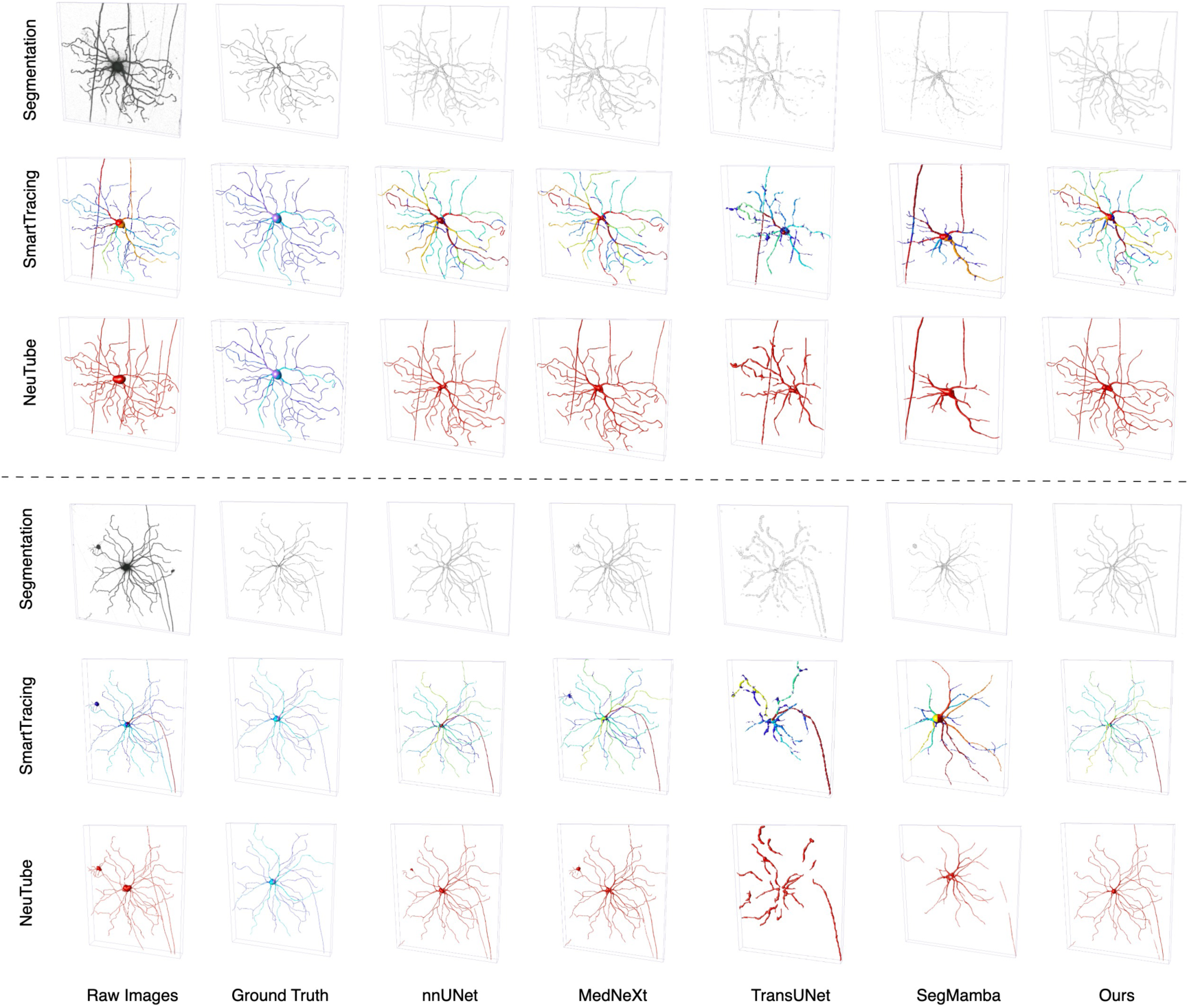
Visualization comparison of the neuron segmentation and tracing performance across different models on the Mouse dataset. The figure shows two representative cases. For each case, the top row presents the raw image and segmentation outputs from different models, followed by two rows showing the corresponding neuron tracing results generated by SmartTracing and NeuTube. Zoom-in views highlighting fine-grained segmentation details are shown in Figure 18.

**Fig. 15.**
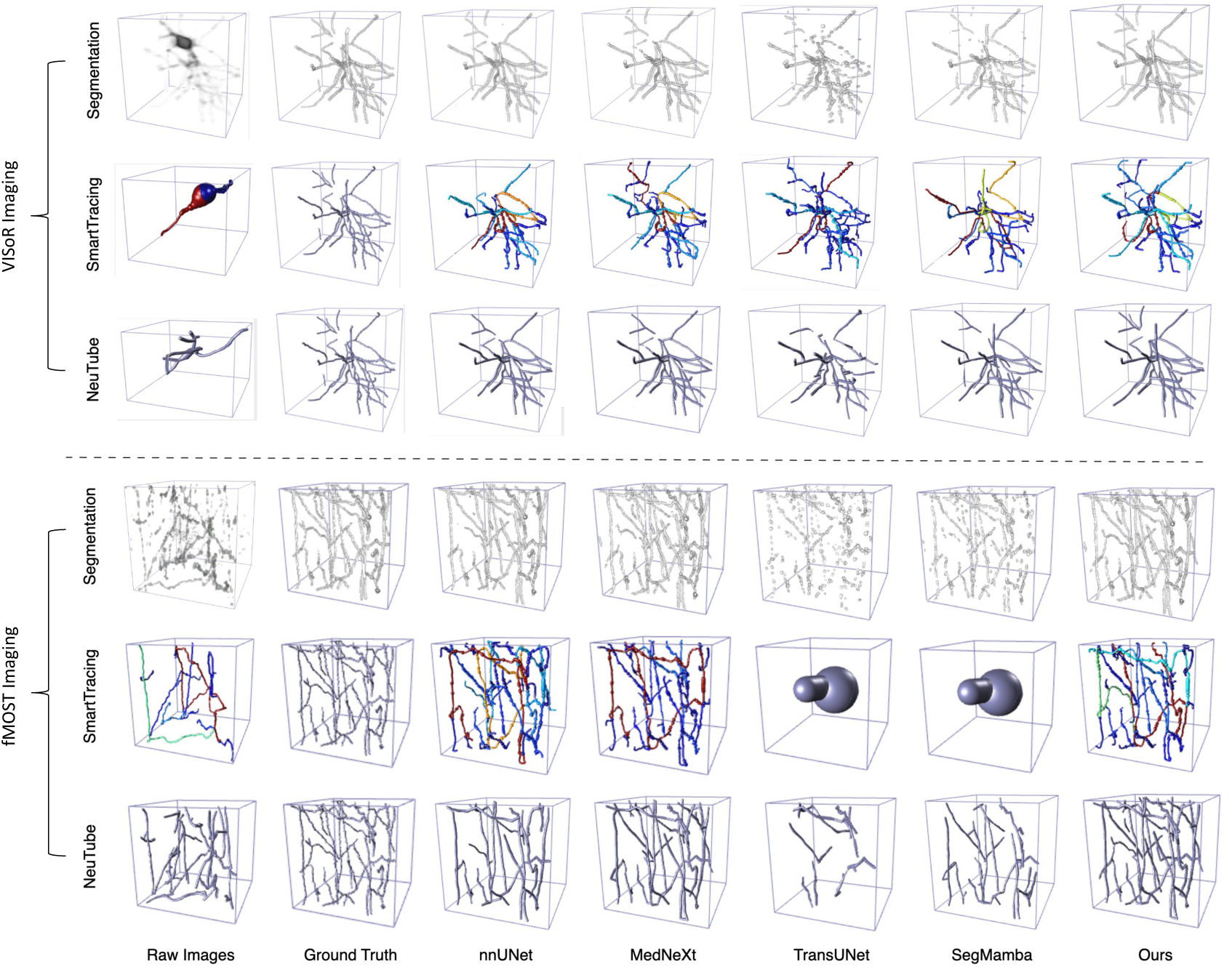
Visualization comparison of the neuron segmentation and tracing performance across different models on the NeuroFly dataset. Two representative 3D volumes are shown: the top block corresponds to VISoR imaging and the bottom block to fMOST imaging. For each case, the top row presents the raw image and segmentation outputs from different models, followed by two rows showing the corresponding neuron tracing results generated by SmartTracing and NeuTube.

**Fig. 16.**
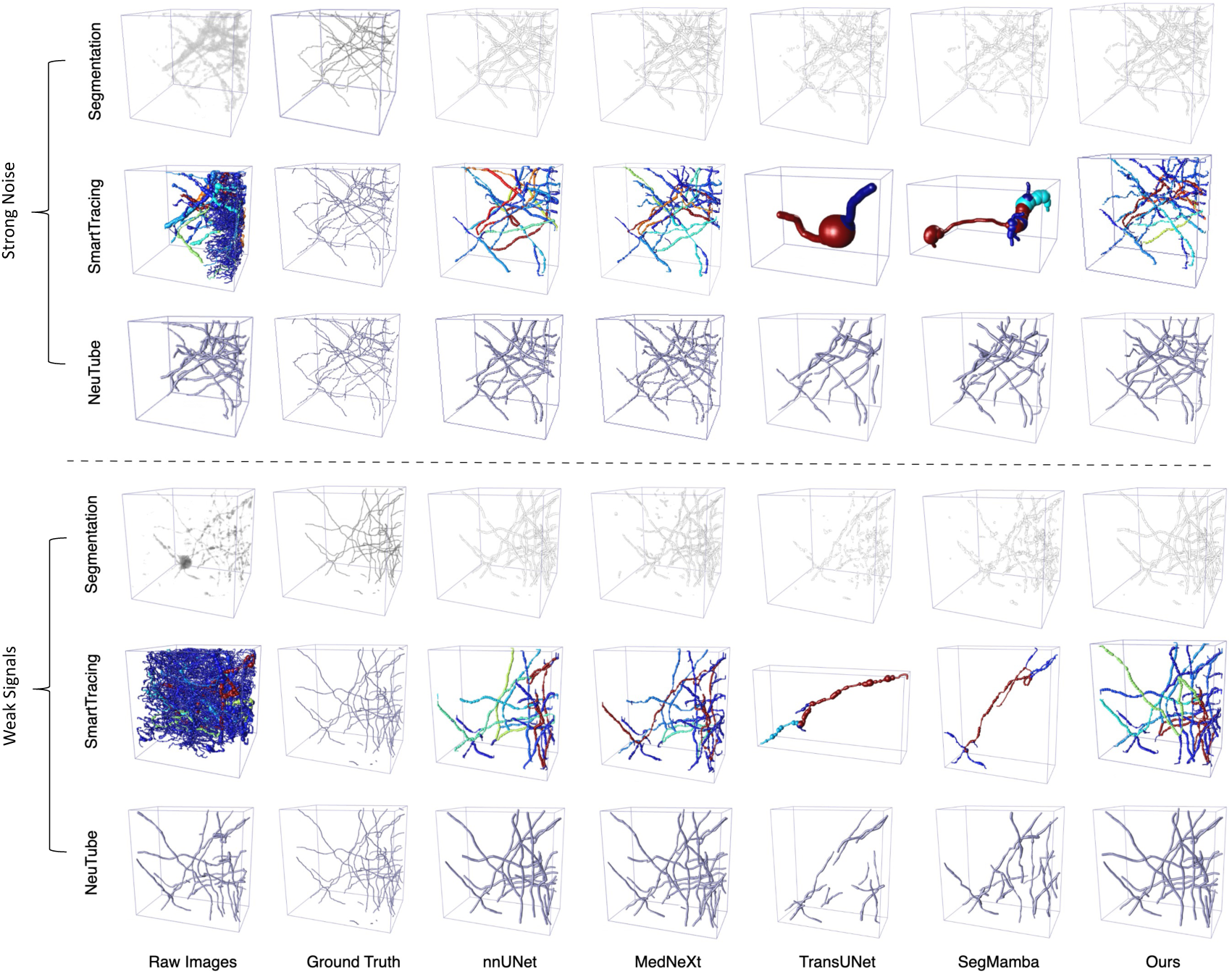
Visualization comparison of neuron segmentation and tracing performance across different models on the CWMBS dataset. Two representative 3D volumes are shown: the top block corresponds to a sample with strong background noise, and the bottom block corresponds to a sample with weak fluorescence signals. For each case, the top row presents the raw image and segmentation outputs from different models, followed by two rows showing the corresponding neuron tracing results generated by SmartTracing and NeuTube.

**Fig. 17.**
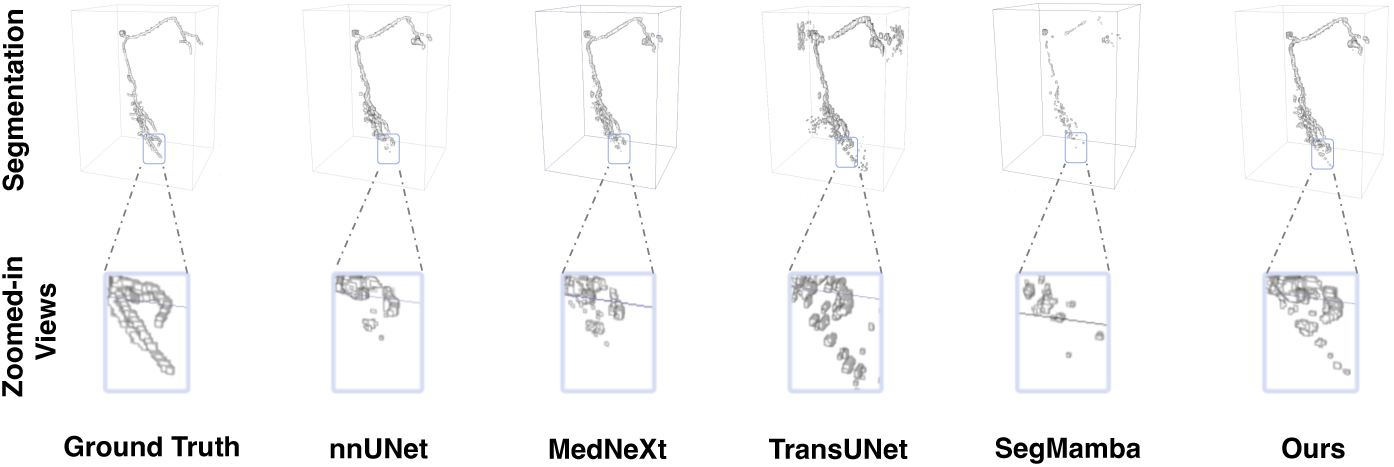
The local details of the segmentation results across different methods on the Drosophila dataset. The top row shows the segmented neuron by different methods, while the bottom row presents zoomed-in views of complex branches. Our method demonstrates improved topological coherence and better preservation of branching patterns compared to others.

**Fig. 18.**
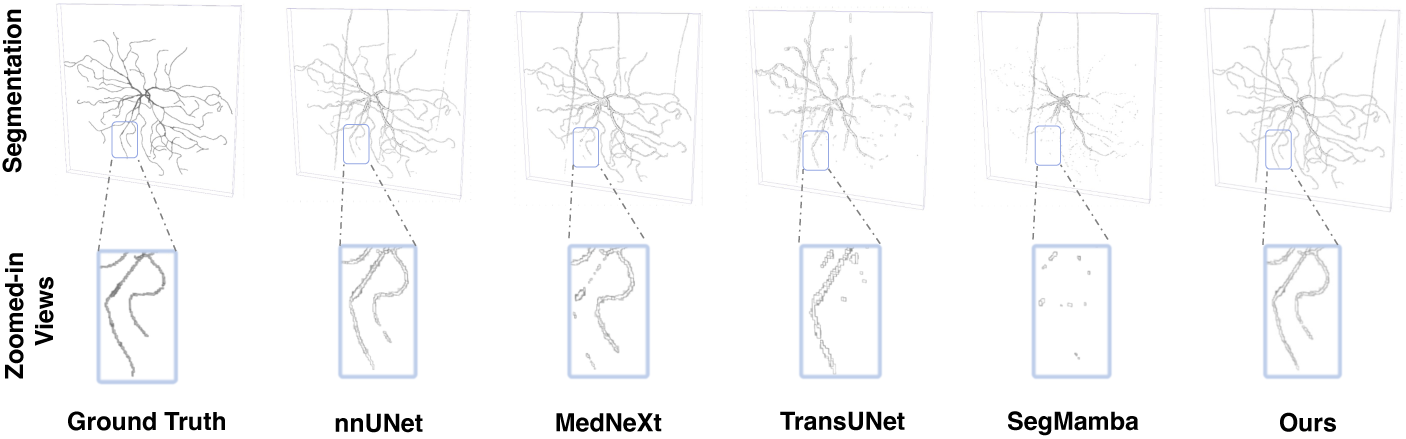
The local details of the segmentation results across different methods on the Mouse dataset. The top row shows the segmented neuron by different methods, while the bottom row presents zoomed-in views of complex branches. Our method demonstrates more continuous and complete structures.

### 4.6 Ablation study

We conduct a comprehensive ablation study on both the Drosophila and Mouse datasets to evaluate the contribution of each component in the proposed DMAC framework. As summarized in Table 1, we systematically investigate four key factors: the dynamic rotation mechanism, the DMAC module, the multi-view strategy, and deep supervision. We report performance on two tracing methods (SmartTracing and NeuTube) using three standard metrics: ESA, DSA, and PDS.

**Table 1.** Ablation study of the proposed DMAC framework on the Drosophila and Mouse datasets.

| Dataset | Variants | Tubular Conv | Rotation | Multi-view | DeepSup | Inference Time (h) | SmartTracing |  |  | NeuTube |  |  |
| --- | --- | --- | --- | --- | --- | --- | --- | --- | --- | --- | --- | --- |
|  |  |  |  |  |  |  | ESA↓ | DSA↓ | PDS↓ | ESA↓ | DSA↓ | PDS↓ |
| Drosophila | Variant 1 (Ours) | ✓ | 2 | ✓ | ✓ | 0.39 | **1.42** | **4.20** | **0.18** | 1.80 | **4.38** | **0.20** |
|  | Variant 2 | ✓ | 1 | ✓ | ✓ | 0.39 | 1.49 | 4.27 | 0.19 | 1.86 | 4.60 | 0.21 |
|  | Variant 3 | ✓ | – | ✓ | ✓ | 0.36 | 1.47 | 4.27 | **0.18** | **1.74** | 4.64 | 0.22 |
|  | Variant 4 | ✓ | 2 | – | ✓ | 0.27 | 1.50 | 4.29 | 0.20 | 1.89 | 4.52 | 0.22 |
|  | Variant 5 | – | – | – | ✓ | 0.17 | 1.80 | 4.43 | 0.23 | 1.95 | 4.82 | 0.25 |
|  | Variant 6 | ✓ | 2 | ✓ | – | 0.39 | 1.47 | 4.26 | 0.20 | 1.83 | 4.45 | 0.21 |
| Mouse | Variant 1 (Ours) | ✓ | 2 | ✓ | ✓ | 1.95 | **2.51** | **7.08** | **0.20** | **4.41** | **14.38** | 0.21 |
|  | Variant 2 | ✓ | 1 | ✓ | ✓ | 1.95 | 2.62 | 7.21 | 0.22 | 4.65 | 14.71 | 0.22 |
|  | Variant 3 | ✓ | – | ✓ | ✓ | 1.82 | 2.65 | 7.26 | 0.21 | 4.60 | 14.79 | **0.20** |
|  | Variant 4 | ✓ | 2 | – | ✓ | 1.33 | 2.70 | 7.34 | 0.22 | 4.67 | 14.83 | 0.23 |
|  | Variant 5 | – | – | – | ✓ | 0.82 | 3.97 | 8.78 | 0.25 | 5.94 | 16.06 | 0.24 |
|  | Variant 6 | ✓ | 2 | ✓ | – | 1.95 | 2.57 | 7.17 | 0.21 | 4.50 | 14.55 | 0.21 |

We evaluate the contributions of key components: tubular convolution, rotation mechanism, multi-view integration, and deep supervision. “Inference Time (h)” denotes the average inference time per test image. Best results are highlighted in bold. Rotation: 2 = two rotation angles, and 1 = single rotation angle.

#### Effectiveness of dynamic rotation

We first examine the impact of the dynamic rotation mechanism by comparing Variants 1, 2, and 3. Variant 1 employs two learnable rotation angles (elevation and azimuth), while Variant 2 uses only one angle, and Variant 3 removes the rotation entirely. The performance consistently degrades as rotation capacity is reduced or removed, with Variant 1 achieving the best reconstruction accuracy across ESA, DSA, and PDS on both datasets. These findings highlight the importance of full 3D orientation adaptability in modeling directionally diverse neuron trajectories.

#### Effectiveness of DMAC

To assess the overall contribution of the DMAC design, we compare the full framework (Variant 1) with a baseline in which all DMAC modules are replaced by standard 3D convolutions (Variant 5). As shown in Table 1, this substitution leads to significant performance degradation across all evaluation metrics. For instance, on the Drosophila dataset with SmartTracing, ESA increases from 1.42 to 1.80, DSA rises from 4.20 to 4.43, and PDS increases from 0.18 to 0.23. Similar degradations are observed on the Mouse dataset. This ablation clearly demonstrates that the proposed rotated tubular formulation is more effective for capturing filamentous and directionally heterogeneous neuronal structures than conventional axis-aligned convolutions.

#### Effectiveness of multi-view strategy

We further evaluate the impact of the multi-view integration strategy by comparing Variant 1 with Variant 4. When multi-view fusion is disabled, the performance consistently deteriorates on all metrics (e.g., ESA increases from 1.42 to 1.50 on Drosophila/SmartTracing). This confirms that integrating features across multiple orientations strengthens feature representation by aggregating complementary directional cues.

#### Effectiveness of deep supervision

We assess the benefit of deep supervision by comparing Variant 1 and Variant 6. Disabling deep supervision (Variant 6) results in inferior performance (e.g., higher DSA and PDS values), indicating that deep supervision effectively facilitates training and improves structural learning across scales.

## 5 Discussion

The proposed DMAC framework achieves consistent improvements in both segmentation and reconstruction tasks across all four benchmark datasets. Its superior performance can be explained by three tightly coupled design elements. First, tubular stretching reshapes the convolution receptive field into a tubular shape to better match the elongated geometry of neuronal fibers. This enables convolution kernel more effective for feature extraction along slender structures while reducing background interference. Second, dynamic rotation aligns the tubular kernel with the dominant local orientation, ensuring that the sampling pattern follows tortuous or oblique neurite trajectories rather than being restricted to fixed axes. Third, the cumulative deformation mechanism further adapts the kernel to capture subtle local curvature and structural variations while preserving spatial continuity along the neurite path. Together, these components are especially beneficial when modeling mesoscaled neuronal morphology, which features extensive arborization and diverse orientations.

The ablation study confirms the contribution of each component. Removing either the tubular convolution, rotation mechanism, or multi-view aggregation consistently degrades performance, indicating that each module plays a complementary role. In particular, multi-view aggregation proves essential for integrating anatomical context from orthogonal directions, while deep supervision improves training and multi-scale segmentation consistency. The trade-off between accuracy and inference time observed in our experiments is largely due to the use of a small patch size and multi-view processing. However, this choice is intentional, as finer patch resolution and richer directional context are critical for accurately capturing intricate neuronal morphologies. In practice, the computational overhead is acceptable for large-scale connectomics applications, where structural fidelity is prioritized over marginal runtime gains.

While DMAC is tailored for filamentous neuronal morphology, this specialization may limit its direct applicability to highly non-tubular or irregular structures. Nevertheless, this morphological bias is precisely what enables the method to excel in preserving neuronal topology and continuity, making it well suited for connectomics and other domains requiring high-fidelity reconstruction of elongated structures.

Overall, the combination of morphology-aware receptive field design, orientation adaptation, and fine-scale deformation control explains the consistent improvements observed in both segmentation metrics (F1, HD95) and reconstruction metrics (ESA, DSA, PDS) across datasets. The framework offers a principled and effective approach to bridging the gap between convolutional feature learning and the unique structural properties of neuronal morphology.

## 6 Conclusion

We presented Dynamic Morph-Aware Convolution (DMAC), a novel segmentation framework that integrates both structural and directional adaptability into the convolutional process to better model the morphological complexity of neuronal structures. By reshaping standard 3D convolution kernels into elongated, deformable tubular forms and dynamically rotating them to align with local neuronal trajectories, DMAC equips the model with structural sensitivity and 3D directional adaptivity tailored to filamentous and variably oriented neuron morphology. These designs improve segmentation continuity and topological fidelity across diverse neuronal geometries. Our extensive evaluations on four mesoscaled neuron reconstruction datasets demonstrate that DMAC consistently outperforms state-of-the-art approaches in both segmentation accuracy and reconstruction quality.

